# Historical squid biomass increase is not explained by rising temperature but rather by loss of top predators

**DOI:** 10.64898/2026.08.30.748117

**Authors:** Rémy Denéchère, P. Daniël van Denderen, Ken H. Andersen

## Abstract

Squid abundance has been reported to increase globally between 1970 and 2010. This increase has been hypothesized to result from two primary factors: the loss of top predators due to overfishing and rising ocean temperatures. The decline in apex predators may lead to the expansion of squid populations either through reduced predation pressure or diminished competition with juvenile predators. Concurrently, increased temperatures could enhance the somatic growth rates of squid, thereby accelerating their population growth. However, empirically disentangling the impacts of predator loss and temperature on squid biomass remains challenging, especially in a food-web context. In this study, we used a size- and trait-based model of upper trophic levels that resolves the ecosystem structure – biomass and trophic interactions of fish and squid – for varying depth, temperature, and secondary production, to investigate two hypotheses of the historical expansion of squid, i.e., the effects of predator depletion from fishing and rising temperatures on squid biomass. Our model reveals that intensified fishing of squid predators – specifically large demersal fish in shelf systems and large pelagic fish in open oceans – leads to a slight increase in squid biomass. Conversely, elevated temperatures are associated with a decline in squid biomass. This temperature-driven reduction in biomass is attributed to an increased metabolism of squids beyond the available food supply. If historic overfishing on large marine predators continues to be curtailed, we expect a corresponding reduction in global squid biomass and fisheries potential, which could be further exacerbated by rising temperatures.

## 1 Introduction

Squid are a valuable global and local economic resource, constituting approximately 4% of the global marine fisheries landings (Arkhipkin et al., 2015; Rodhouse, 2005). In addition to their direct economic value, squid serve as prey for large predatory fish, indirectly supporting commercial fisheries (Hunsicker et al., 2010), and are a significant food source for tooth-whales (Garibaldi and Podestà, 2014). They also have the potential to sequester carbon more effectively than fish due to their fast lifestyle and reproductive strategy (Ottmann et al., 2024), demonstrating their substantial impact on ecosystem structure and services.

The ecological significance of squid rests on their distinctive life-history traits. Their rapid growth demands voracious feeding to meet their metabolic requirements (Rodhouse et al., 1998, Chap. 13). Despite their relatively large size, squid have a remarkably short lifespan, as demonstrated by *Doscidicus gigas*, which typically ranges from 10 to 100 kg in about 1 to 1.5 years,(Goicochea-Vigo et al., 2019). Their short lifetime is supported by their distinctive fast somatic growth, about 5 times faster than teleost fish, and semelparous reproductive strategy (Denéchère et al., 2024; Laptikhovsky et al., 2019). Squid are often described as opportunistic species, capable of rapid expansion when environmental conditions are favorable (Boyle and Boletzky, 1996).

Evidence suggests that cephalopod fishery landings have increased significantly during the latter half of the 20th century (Caddy and Rodhouse, 1998). A more recent global study by Doubleday et al. (2016) reported a rise in the abundance of squid and other cephalopods across multiple regions. The authors proposed two key drivers for this trend: the loss of top predators due to overfishing and increased sea temperatures from global climate change. In recent years, these hypotheses have been widely cited in the literature (Ospina-Alvarez et al., 2022; Rosa et al., 2019; Gleadall et al., 2018; Arkhipkin et al., 2021; Purkayastha and Furuya, 2026), at times without acknowledging their speculative nature (Guerreiro et al., 2025; Pauly and Froese, 2021; Golikov et al., 2025). However, the proposed explanations for the historical increase in squid abundance are speculative, and the underlying mechanisms remain unclear.

Here, we investigate the effect of both temperature and fishing of large predatory fish as two hypotheses for the historical expansion of squid described by Doubleday et al. (2016). The hypothesis that predator depletion promotes squid proliferation is conceptually straightforward. Industrial fishing has historically reduced the biomass of large marine predators (Mathiesen, 2015; Pauly et al., 1998). A decline in top predators should relax predation mortality on squid and reduce interspecific competition. Empirical evidence provides partial support for this mechanism. For instance, Caddy and Rodhouse (1998) suggest that overexploitation of groundfish stocks has enhanced squid and octopus populations. Similarly, Vecchione et al. (2009) documented an increase in octopus abundance off the Antarctic Peninsula coincident with overfishing. These studies suggest that over-fishing may facilitate cephalopod expansion; however, the authors emphasize the need for further testing this hypothesis across ecosystems. Predator depletion may also restructure trophic interactions into a cephalopod-dominated system with potential intensification of intra-specific competition and cannibalism. Such dynamics are characteristic of trophic cascades documented in heavily exploited systems, including the North Sea (Daan et al., 2005) and the Northwest Atlantic (Frank et al., 2005). Nonetheless, the net effect of predator depletion is largely expected to benefit squid populations, allowing their expansion in regions where top predators have been heavily fished.

Temperature is a known factor impacting the physiology and life cycle of squid, affecting their embryonic development (see discussion), metabolism, and reproduction (Pecl and Jackson, 2008). Temperature has a direct effect on somatic growth, respiration, and maximum consumption rates of squid. As temperatures rise, metabolic rates increase, which may result in accelerated growth and higher population growth rates. These results have been widely supported by experimental, field, and theoretical studies (Hatfield et al., 2001; Forsythe, 2004; Denéchère et al., 2022; Villanueva, 2000; Forsythe et al., 2001). The Forshythe hypothesis clearly states that temperature alone is not sufficient to predict growth in nature and must be understood in a wider context, including food assimilation (Forsythe, 2004). More specifically, any temperature-induced increase in growth must only occur under the condition of sufficient food intake. This argument logically arises from a common understanding that an individual’s energy can be viewed as a budget allocated to all physiological processes, including growth (O’DOR, 1987; Lee, 1995; Wells and Clarke, 1996; Kooijman, 2010). Evidence supporting this is limited, but Jackson and Domeier (2003) demonstrated how hydrographic conditions can alter food supplies and thus squid growth. Following a strong El Niño/La Niña effect off the southern California, reduction in food availability can override temperature-enhanced growth. In natural environments, it remains uncertain whether squid can meet the increased metabolic demands associated with higher temperatures, raising concerns that warming may ultimately constrain population growth.

To investigate hypotheses behind the historical expansion of squid, we used the size-based model of high trophic level communities, FEISTY-squid (Denéchère et al., 2024). The FEISTY-squid framework resolves the community structure of several functional groups of fish – small and large pelagic, demersal, and mesopelagic – and squid. It characterizes the energy flow and biomass distribution of fish and squid in a broad range of ecosystems from shelf systems to the open ocean by resolving the vertical distribution and benthopelagic coupling resulting from two environmental drivers: the bottom depth and secondary production. We explored the effect of increasing fishing on squid predators represented by large demersal fish in shelf regions and large pelagic fish in open oceans. We further studied the effect of increasing temperature on squid metabolic rates and its consequences for the biomass of squid and fish. We tested the impact of temperature with several hypotheses on the type of *Q*_10_ and mass-scaling of vital rates of fish and squid.

## 2 Materials & Methods

In the following, we provide an overview of the functioning of the FEISTY-squid framework, with particular attention to the two central aspects of this paper: fishing and temperature. The detailed formulation of the FEISTY-squid model, including the underlying equations and sensitivity analyses, can be found in Denéchère et al. (2024).

### 2.1 FEISTY-squid framework

FEISTY-squid is a size- and trait-based model based on the fish model from van Denderen et al. (2021). The model groups high-trophic levels into five functional types – small and large pelagic fish, demersal fish, mesopelagic fish, and squid – each characterised by its asymptotic mass *M*, individual growth rate *h*, and vertical feeding habitat. Squid are parametrised with a growth rate coefficient five times higher than fish, reflecting their fast life history (Denéchère et al., 2024). Two resource pools support the community: zooplankton and benthic invertebrates, imposing competition between and within functional groups.

All processes operate at the individual level, defined by body size and modulated by temperature. The core is a Dynamic Energy Budget (DEB) model where prey are consumed based on encounter rates, then assimilated and respired, and the remaining available energy *v*_*i*_ (*m*)is divided between somatic growth and reproduction according to maturity (Fig. 1A). The governing equation is a discretization of the McKendric-von Foerster equation (De Roos et al., 2008), which describes the change of biomass in size class *i* due to fluxes between neighbouring classes, reproduction, and mortality (see Methods section from Zhao et al. (2024)):

**Figure 1.**
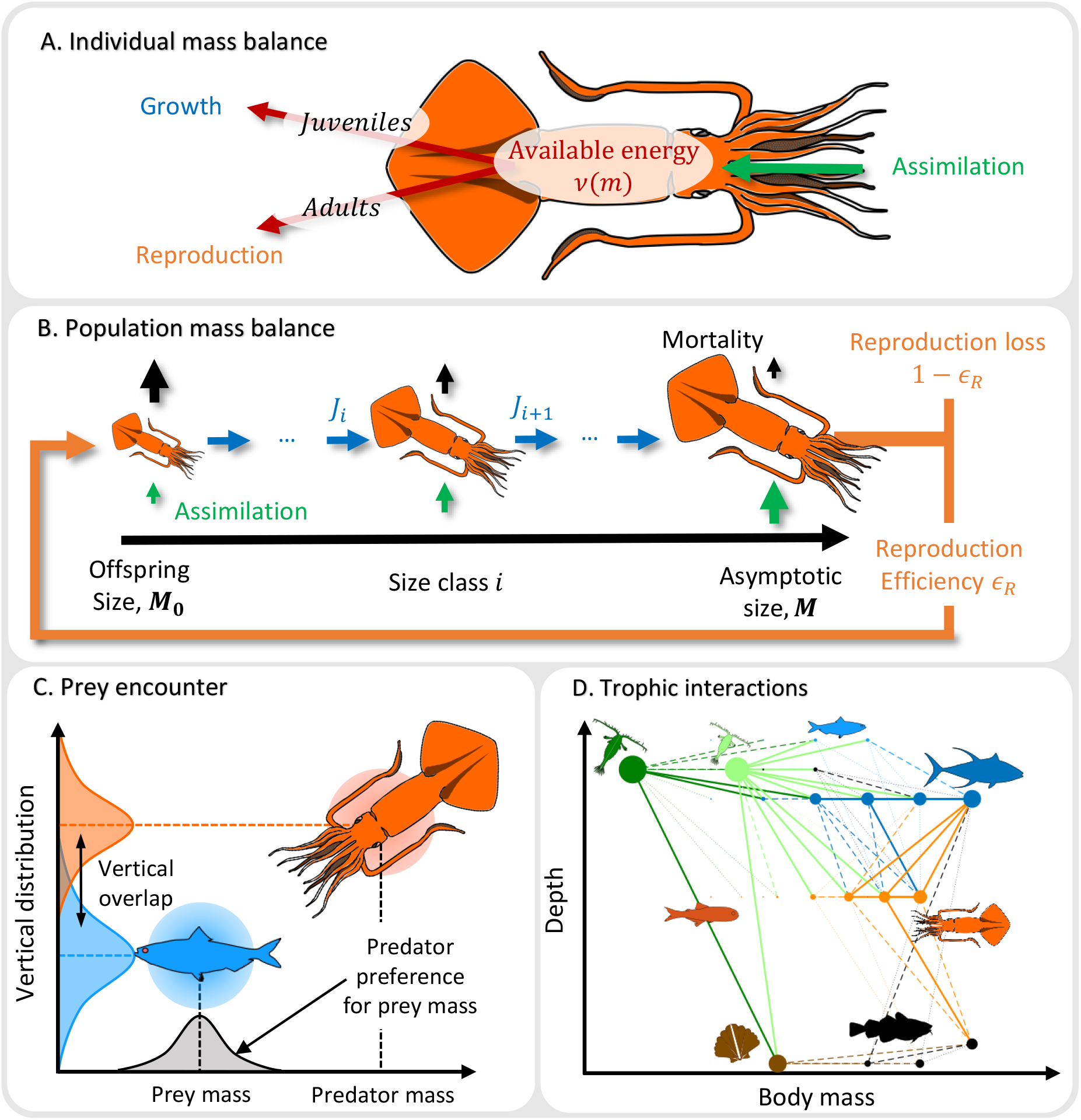
Squid in the FEISTY model. (A) Dynamic Energy Budget model of squid. (B) Population mass balance. (C) Prey encounter. (D) Trophic interaction within FEISTY-Squid. The width of the dots represents the biomass of the size class. The colors indicate the functional groups: small and large zooplankton (dark and light green, respectively), benthic resource (brown), demersal fish (black), small and large pelagic fish (light and dark blue, respectively), mesopelagic fish (red), and squid (orange). The line thickness represents the intensity of the flux of prey eaten per mass of predator, from low (dotted lines), medium (dashed lines), to high (plain lines).

**Figure 2.**
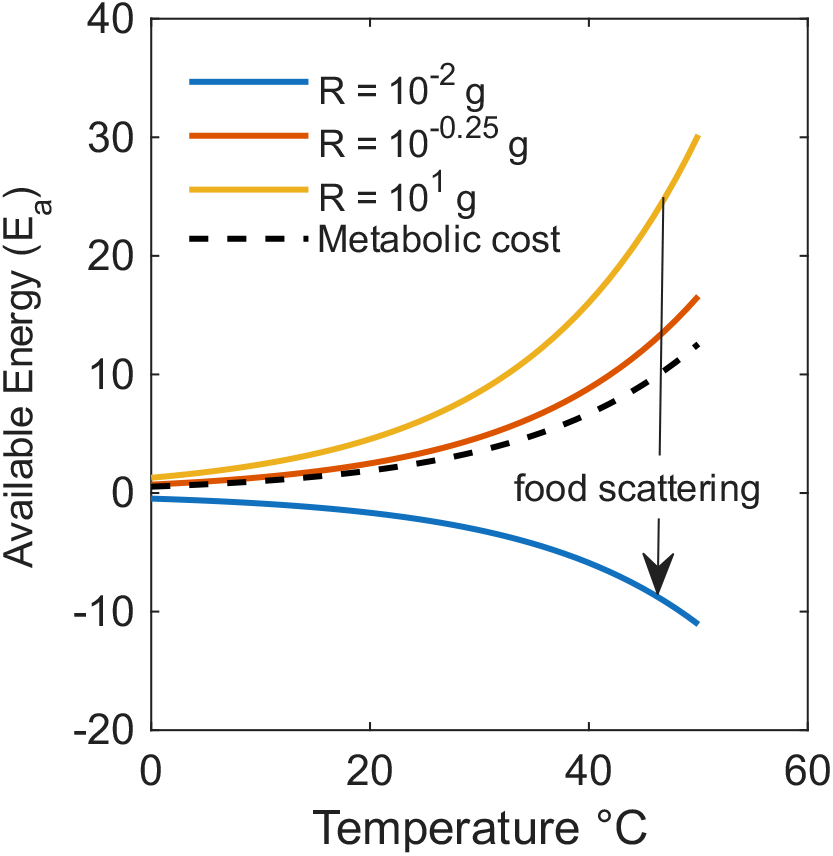
Available energy for increasing temperature and three levels of decreasing resources. Increasing metabolic cost with temperature is represented by the black dashed line.

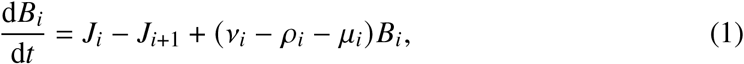

with *B*_*i*_ the biomass (units of biomass) of the *i*th size class, *J*_*i*_ and *J*_*i* 1_ are the fluxes of biomass in and out of the size class *i* (units of biomass per time), *ρ*_*i*_ is the biomass-specific reproduction rate (units of per time), and *µ*_*i*_ is the biomass-specific mortality rate (units of per time) and *v*_*i*_ represents the biomass-specific available energy after accounting for basal metabolism, which fuels growth and reproduction in the adult size classes (Fig. 1A & B).

Community structure is defined as the biomass of each functional group and trophic relationships that emerge from size-based predation preferences, where large predators feed on smaller prey and share vertical distributions in the water column (Fig. 1C & D).

### 2.2 Fishing on large predators

Mortality *µ*_*i*_ (*m*) in the FEISTY framework results from two primary sources: predation by other groups *µ*_P,*i*_ (*m*) and fishing *µ*_F,*i*_ (*m*). This is represented as:

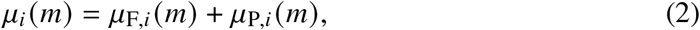

where *µ*_P,*i*_ (*m*) reflects the predation mortality, calculated based on the vertical overlap of prey and predators and size (van Denderen et al., 2021; Zhao et al., 2024). We impose fishing mortality *µ*_F,*i*_ (*m*) on squid predators in FEISTY, specifically demersal fish in the shelf system and large pelagic fish in the open ocean. The fishing mortality rate for squid and small pelagic fish is kept constant at 0.1 yr^*−*1^. We assume that fishing is mainly due to trawling fisheries for which the fishing mortality function follows a logistic function with size as described by Myers and Hoenig (1997) and Andersen (2019):

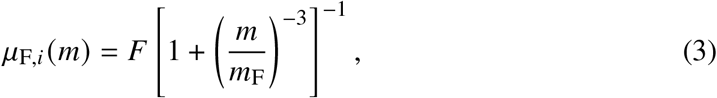

where *m* is the body mass of the size class *i, m*_F_ is the size-selectivity for fishing (in grams wet weight) and represents the size at which the selectivity equals 50% of the fishing intensity, and *F* is the fishing intensity (per year). We assume the size-selectivity for fishing to be 5% of the adult size (*m*_F_ = 0.05*M*) following Andersen (2019).

### 2.3 Temperature

The FEISTY-squid model assumes that temperature affects the maximum consumption rate *C*_max_, the searching rate *V* (foraging activity), and the basal metabolism *M*_c_. *C*_max_ and *V* scale with individual mass with exponents *−*0.25 and *−*0.2, respectively, and are two components of individual consumption in FEISTY (Eq. 4 & 5). The basal metabolism represents the respiration cost to maintain individual mass and scales with a *k* = *−*0.1 exponent with body size (Eq. 6). The effect of temperature is described as *Q*_10_ factors, such that the above physiological process can be written as resulting from both size and temperature:

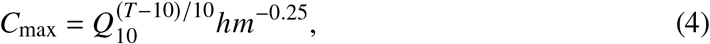

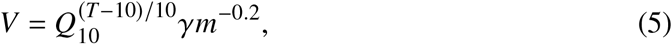

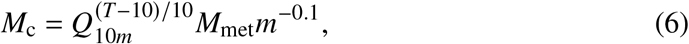

where *h* is the coefficient of maximum consumption, *γ* is the coefficient for searching rate and *M*_met_ the coefficient for metabolic cost. The temperature *T* (°C) represents the average experienced by a functional group. Therefore, *T* is assumed to be constant with depth, and the vertical position of the functional group does not change the experienced temperature. Note that the coefficient for maximum consumption *h* is interpreted as an individual’s potential for somatic growth and links to the metabolic cost such that *M*_met_ = 0.2*h*. In the FEISTY-squid model, we assume that squid have a higher *h* compared to fish (100 g^*n*^ yr^*−*1^ for squid versus 20 g^*n*^ yr^*−*1^ for fish), resulting in squid exhibiting elevated metabolic demands and greater food requirements to sustain their population (see Denéchère et al. (2024)).

Equations 4 to 6 define the energy available for growth and reproduction (*E*_a_) as:

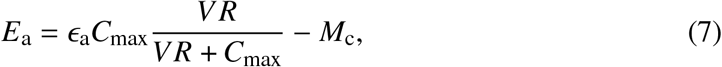

where *ε*_a_ is the assimilation efficiency, and *R* the resource concentration in the environment. This equation establishes a balance between consumption and metabolic cost in the sense that consumption has to exceed metabolic cost for the organism to grow or reproduce. As temperature increases, it leads to higher foraging activities and greater food consumption, but it also results in increased metabolic costs (Fig.2 black dashed line). Note that in our model, temperature affects reproduction indirectly through its influence on food consumption and metabolic cost.

The effects of temperature on squid are tested by varying temperature by two degrees Celsius from the “baseline” parameters (10°degrees Celsius) and re-running the model to equilibrium conditions. This means that the model does not explicitly resolve the temperature performance for individual species and their thermal niche, but rather the consequence of the temperature change on the dynamics of squid biomass over longer time scales, scales where migration and evolution have adjusted local ecosystems. We test two key uncertainties regarding the effect of temperature on squids: 1) does temperature affect the three parameters in the energy budget (*V*, *C*_max_, and *M*_c_) similarly; and 2) do squid have a similar *Q*_10_ to fish? These tests are done by adjusting the effects of temperature on the clearance rate and maximum consumption rate (*Q*_10_) and metabolic rate (*Q*_10*m*_), as well as adjusting *Q*_10_ of fish and squid independently (see discussion and Supplement B and C).

### 2.4 Modelling experiment

We examine the effect of temperature and fishing in a shelf system (50 m depth) and an open ocean (2000 m depth) system. We parameterise the model with high zooplankton productivity to ensure that squid are present in the system (see Denéchère et al. (2024)). In the shelf system, FEISTY exhibits strong benthopelagic coupling with interactions between demersal fish, squid, and pelagic fish. In the shelf system, squid are eaten mainly by large demersal fish (Fig. 1A). There is no benthopelagic coupling in the open ocean system, and most interactions occur in the pelagic layer and between the pelagic and mesopelagic zones. Squid are mainly eaten by large pelagic fish in open oceans (Fig. 1B). “All simulations were run in MATLAB R2024. Code is available at: https://github.com/RemyDenechere/Squid_biomass_decreases_with_temperature.git

## 3 Results

### 3.1 Fishing

In both the shelf and open ocean the increased fishing on top predators is associated with a slight increase in squid biomass (from 8.5 to 15 g m^*−*2^ in the shelf system Fig. 3A; and from 11 to 22 g m^*−*2^ in the open ocean Fig. 3B). The increase in squid biomass is small compared to the decline of top predators; from about 64 to 7.5 g m^*−*2^ for demersal in the shelf system (black line; Fig. 3A); and from about 82 to 0.0 g m^*−*2^ for large pelagic in the open ocean (thick dark-blue line; Fig. 3B). This large decrease of top predators leads to a decline in total biomass in the ecosystem.

**Figure 3.**
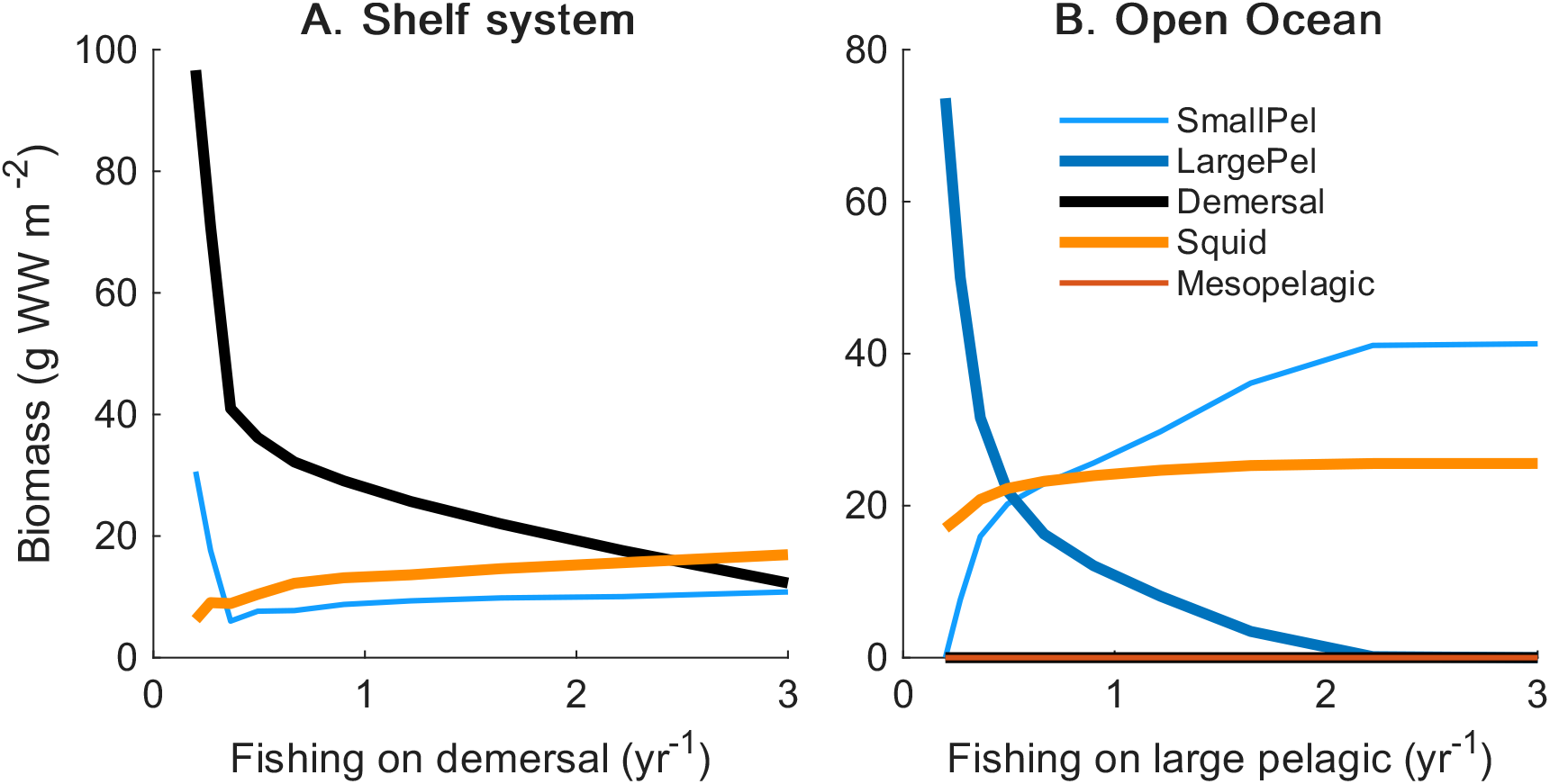
Effect of fishing intensity *F* on squid predators – demersal in a shelf system (50 m depth) and large pelagic in an open ocean (2000 m depth) – on Biomass (lines) and on squid proportion (shaded areas). Fishing intensity is constant and equal to 0.1 yr^*−*1^ for the other groups.

The fraction of food content in squid stomachs remains unchanged despite the increase in squid biomass (Fig. 4A & B), suggesting that the competition for food experienced by squid does not intensify with increased fishing pressure in our model. However, squid shift the predominant part of their diet from juvenile large pelagic fish to smaller pelagic fish as fishing mortality on large pelagic fish increases. Interestingly, despite the reduction in top predators, the level of predation experienced by squid increases (Fig. 4C). We attribute this to increased cannibalism, evident from higher squid proportions in stomach contents under high fishing (Fig. 4A & B, orange).

**Figure 4.**
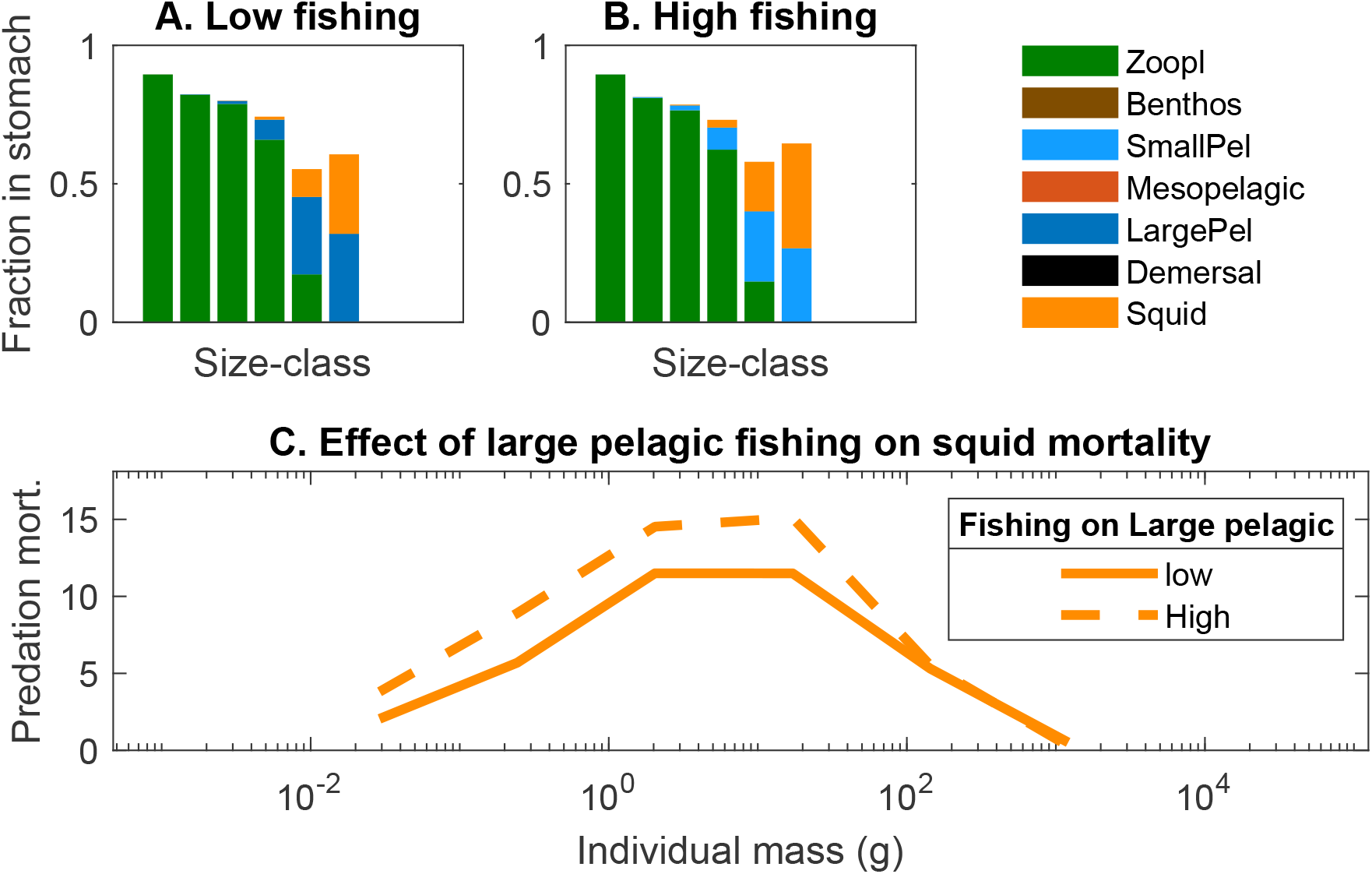
Effect of large pelagic fishing on squid feeding level in an open ocean system (2000 m depth). Top panels (A and B) represent the fraction of food in squid stomachs (consumption divided by maximum consumption *C*_max_). Panel C shows the predation mortality experienced by squid in two cases: low and high fishing mortality on large pelagic (squid’s predators) are characterised by fishing intensities of 0.1 and 3 yr^*−*1^, respectively.

### 3.2 Temperature

In both the shelf and open ocean systems, a temperature increase of 2°C decreases squid biomass and increases fish biomass (Fig. 5). At low zooplankton productivity, squid are absent from the system and an increase of 2 °C results in lower fish biomass (Fig. 5; black dashed lines; ≤ 25 g m^*−*2^ yr^*−*1^). At higher productivities, an increase of 2 °C positively affects fish biomass, resulting in a lower squid proportion in the system.

**Figure 5.**
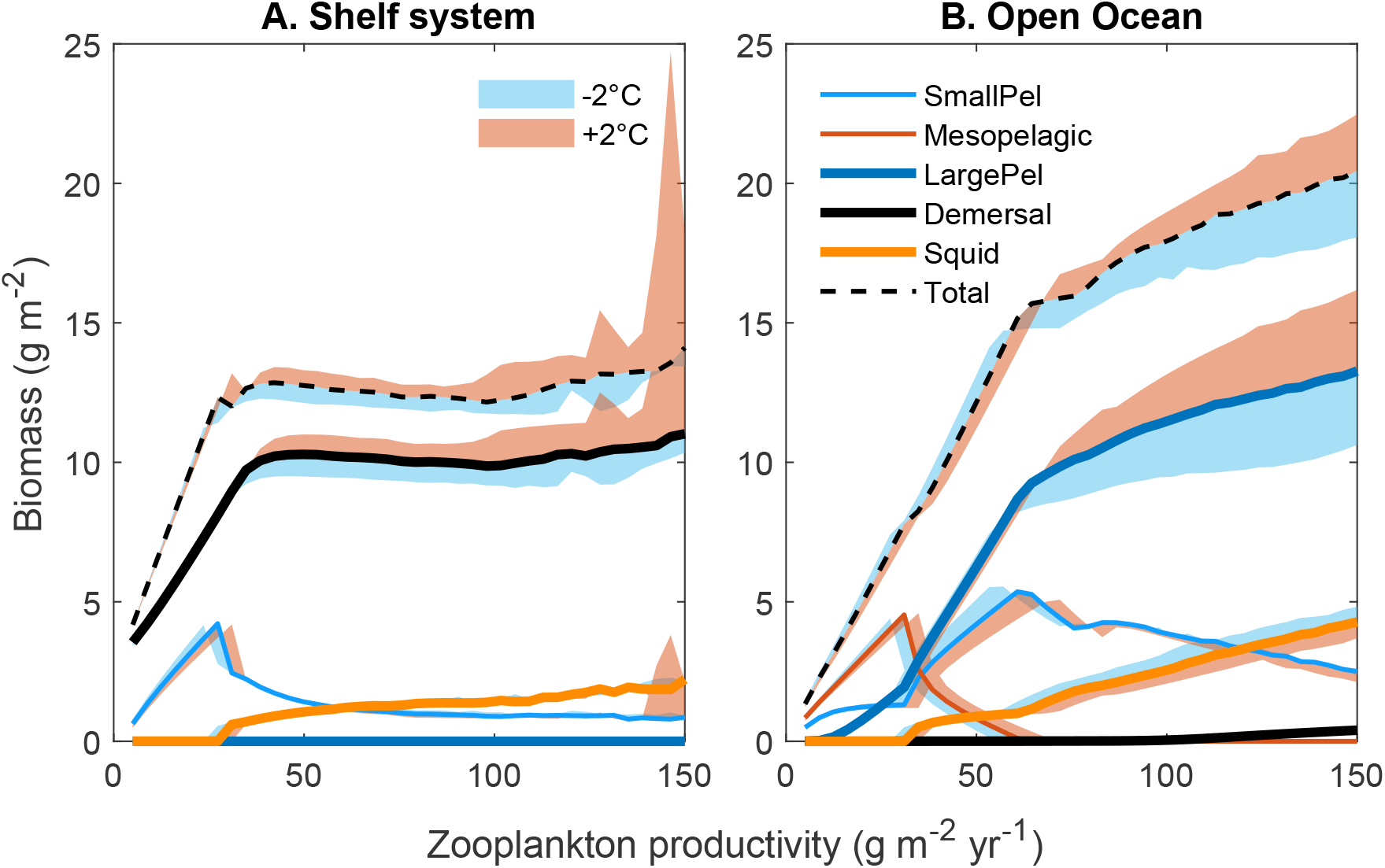
Simulation of the biomass in the FEISTY-squid model for increasing zooplankton productivity for three temperature scenarios: at the standard 10 °C (lines), at the standard *−*2 °C (blue shaded area) or +2 °C (red shaded area) for a shelf (50 m depth) and a deep (2000 m depth).

## 4 Discussion

The loss of top predators leads to a net increase in squid abundance, though this increase is minor compared to the significant decline in predatory fish biomass. Interestingly, higher temperatures reduce squid’s biomass, indicating that squids cannot compensate for temperature-driven increased metabolic costs by consuming more food (see details below). Our model shows that temperature and predator loss have contrasting effects on squid biomass, suggesting that the historical increase in squid biomass is more likely driven by declines in large squid predators rather than an increase in temperature.

### 4.1 Fishing

Our results indicate that although squid biomass increases with the loss of top predators, the increase is fairly small. This suggests that squid are more regulated by bottom-up (resource concentration) rather than top-down processes (predation).

Bottom-up control is further evident in Fig. 5, where squid biomass scales with secondary production. Previous studies have also shown that squid are controlled by bottom-up processes. First, they exhibit high annual variability that is explained by environmental conditions of temperature and resource availability (Boyle and Boletzky, 1996). Second, squid have a short-time life cycle associated with a fast growth rate and high metabolic costs (Denéchère et al., 2024; Goicochea-Vigo et al., 2019; Arkhipkin and Silvanovich, 1997), and, consequently, a voracious feeding behaviour (Rodhouse et al., 1998, Chap. 13). Third, our work highlights a significant degree of cannibalism in squid populations, a finding that is supported by empirical studies through visual observations of predation and stomach content analyses (Hoving and Robison, 2016; Vovk, 1985; Phillips et al., 2003). Together, these results suggest intra-specific or intra-group regulation rather than top-down effects originating from large predatory fish.

Since squid are mainly regulated by bottom-up processes and their intra-group mortality increases with predator loss, any impact of predator decline on the historical increase in squid abundance (Doubleday et al., 2016; Caddy and Rodhouse, 1998) must stem from indirect effects, such as reduced competition or an increase in smaller and more productive prey. Further, as demonstrated in our model, the loss of top predators enhances the abundance of smaller forage fish, which are more productive and serve as key prey for squid. The effect of predator loss therefore indirectly favors the squid by increasing their available food.

### 4.2 Temperature increases the resource demand

The temperature in the FEISTY-squid increases both the searching rate for food and the metabolic cost of squid (and fish). Our FEISTY-squid model has shown that squid are constrained by food availability (Fig. 5; and Denéchère et al. (2024)), and have little opportunity to further increase consumption with increasing temperature. In nature, these effects may be weakened as squid may be able to adjust their metabolic demand to their resource environment (Clarke, 2003; **?**). This highlights the importance of studying the interplay between temperature and resource availability, which is not yet understood for squid.

Our work only includes changes on metabolic rates and consumption rates of fish and squid due to increasing temperature. Yet, temperature also affects egg/larval development as well as fish and squid resources by changing the physical environment, such as stratification of the water (Behrenfeld et al., 2006) and plankton physiological processes (Allen et al., 2005; Brown et al., 2004; Thomas et al., 2017). The effects of temperature on the physiological processes of plankton are complex (Serra-Pompei et al., 2019) and not examined here.

### 4.3 Squid as first invaders

Our work showed that the historical expansion of squid is not, or only partially, explained by temperature and the loss of top predators, respectively. Then, how to explain this historical expansion? An alternative mechanism that may lead to squid expansion is their invasion potential. There are two conditions for a species to be a good invader: having a fast growth rate and a short life span. Species with a fast population growth rate, i.e., an *r−* type strategy, usually have a low competitive advantage when the resource is limited; however, they can invade rapidly when the conditions are favourable (Pianka, 1970; MacArthur and Levins, 1967). Generally, small species are considered *r−*types, while large species employ a *K−* strategy. Thus, the consideration that squid belong to the *r −*strategy has to be made in regard to other groups of the same size, such as fish. Indeed, squid grow 5 times faster than fish (Denéchère et al., 2024). Therefore, they have a higher maximum population growth rate *r*_max_ than fish (Denéchère et al., 2022). Usually, *r−K* strategies are only considered in relation to the potential growth rate of a population *r*_max_ (Pianka, 1970) but not in relation to maturation time. However, maturation time *t*_mat_ is a fundamental metric to consider in the case of invasion potential as it describes the gap between the first invasion and the next generation. *t*_mat_ cannot be calculated directly from the somatic growth rate but depends on both the size at maturation and the somatic growth of the population (Andersen, 2019). Squid have a very short lifespan of about 1 to 2 years. For instance, *Doscidicus gigas* has an average life span of 1*−*1.5 years (Goicochea-Vigo et al., 2019). As a result, squid clearly exhibit an *r* life history strategy, with high potential for invading new environments.

The invasion potential of squid allows them to invade quickly in environments characterised by high disturbance, i.e., with high variation in environmental conditions. Disturbance and/or variability have potentially increased in ecosystems due to climate change and fishing. For instance, heat waves have increased in frequency in the last century (Oliver et al., 2018). Similarly, the decline of top predators due to fishing may have reduced food-web stability (Rooney et al., 2006). Additionally, fishing practices, such as bottom trawling, have altered benthic shelf systems with changes in community composition and function (Van Denderen, 2015; Van Denderen et al., 2015). Considering this, the increase in squid populations in the last century could be alternatively explained by the ability of squid to quickly respond to increased ecosystem disturbance.

### 4.4 Caveat

We treated temperature as constant over depth. Temperature often decreases with depth. Squid in the open ocean appear to be mesopelagic, i.e., living in deeper and colder layers. Their vertical niche makes them less sensitive to temperature variation at the surface as they only spend a fraction of their time in the pelagic zone. As a result, their sensitivity to surface temperature variation is likely attenuated relative to what our model assumes.

*Q*_10_ values for squid are rare and poorly established. We identified two studies examining *Q*_10_ values for squid, with differing results by size and species. Segawa (1995) reported values from 2.01 to 1.60 for the Oval squid (*Sepioteuthis lessoniana*) between 0.4 and 3.6 g, though these may not extend to larger sizes. Trueblood and Seibel (2013) found a *Q*_10_ of 2.1 for adult jumbo squid (*Dosidicus gigas*). Given this variability and the limited data on temperature effects on specific physiological rates (e.g., clearance rate, maximum consumption, and metabolic rate), we tested scenarios with *Q*_10_ values lower, equal to, and higher than *Q*_10*m*_ for Squid. Our results consistently showed a negative impact of temperature on squid biomass (see Supplementary B). Additionally, we varied *Q*_10_ assumptions between fish and squid, where squid *Q*_10_ values were lower, equal to, or higher than those of fish, while keeping *Q*_10_ = *Q*_10*m*_ within each group. This variation did not significantly affect temperature-driven biomass predictions.

Our model does not account for the effects of temperature on squid embryonic development. Warmer temperatures accelerate embryonic development, which often results in smaller hatchling sizes (Boletzky, 1994; Vidal et al., 2002; Bouchaud, 1991). Smaller hatchling sizes may lead to reduced adult sizes, which could have cascading effects on population dynamics. For instance, smaller offspring are more susceptible to predation, as mortality rates generally decrease exponentially with increasing body size (Andersen, 2019, Chap. 2). Including these effects could enhance the model’s ability to capture temperature-driven changes in squid populations.

## 5 Conclusion

Here, we show that the expansion of squid populations of the last century can be partly explained by a loss of top predators. The effect of an increase of temperature is unlikely to explain squid expansions as squid are mainly regulated by their resource. As a consequence, increasing temperature results in a decreasing squid biomass. Besides fishing effects, squid’s fast life history strategy allows them to expand when conditions are unstable.

