## Supplementary materials for "Historical squid biomass increase is not explained by rising temperature but rather by loss of top predators"

### Supplement A:

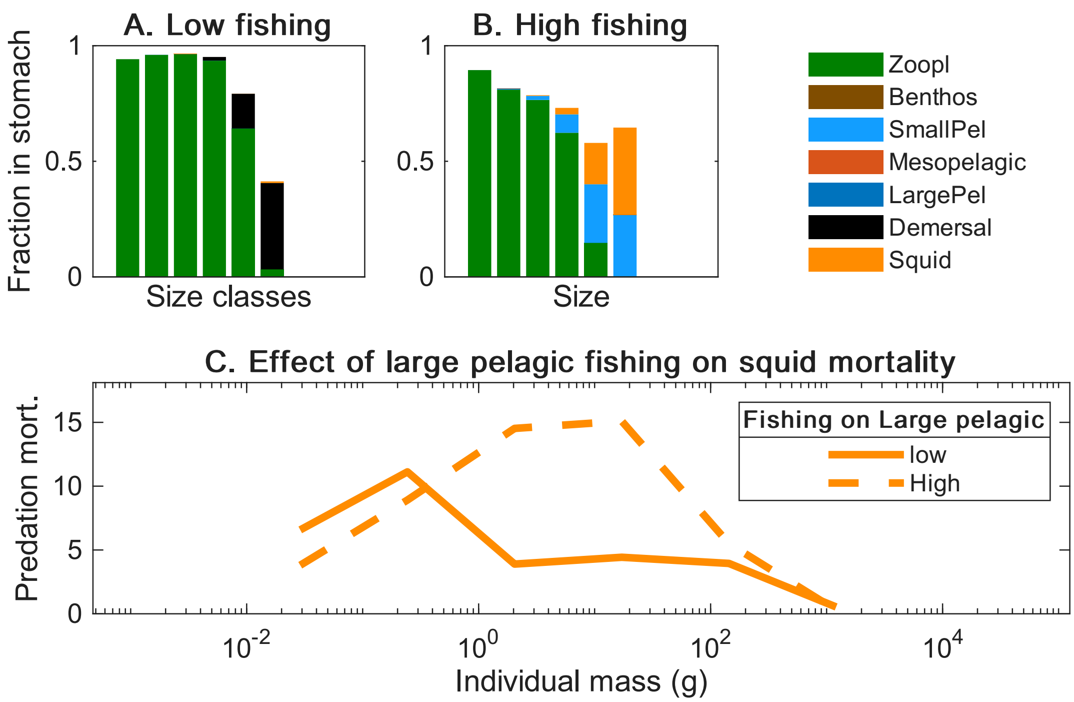


Figure SA1: Effect of large pelagic fishing on squid feeding level in an shallow system (50 m depth). Top panels (A and B) represent the fraction of food in squid stomachs (consumption divided by maximum consumption $C_{\max}$). Panel C shows the predation mortality experienced by squid in two cases: low and high fishing mortality on large pelagic (squid's predators) are characterised by fishing intensities of 0.1 and 3 $yr^{-1}$, respectively.

### Supplement B: Effect of varying $Q_{10}$ assumptions for squid.

We tested three configurations for the effect of temperature on maximum consumption and clearance rates ($Q_{10}$) and metabolic cost ($Q_{10m}$): case 1, where $Q_{10}$ exceeds $Q_{10m}$; case 2, where they are equal; and case 3, where $Q_{10}$ is lower than $Q_{10m}$ (see Table. SB1)

Overall, the biomass of squid decreases with increasing temperature. This happens for each of the above cases. In the shelf ecosystem (Fig. SB1A), the different cases of $Q_{10}$ for squid do not affect the general trend with temperature, and all show a slight decrease in squid biomass and an increase in demersal and small pelagic biomass. In the open ocean (Fig. SB1A), the differences in squid $Q_{10}$ do not affect the squid biomass decrease with temperature. For fish, there is some variation in the slope of biomass decrease with temperature. Demersal and large pelagic biomass increase regardless of the $Q_{10}$. In the case of small pelagic, the biomass is decreasing when squid consumption has a higher sensitivity to temperature than metabolic rate, i.e., in the case where $Q_{10} > Q_{10m}$. Conversely, their biomass increases when squid consumption increases less than the metabolic rate with temperature, i.e., in the case where $Q_{10} < Q_{10m}$.

Table SB1: Cases of $Q_{10}$ values for squid tested for temperature simulations. $Q_{10}$ values of fish are also indicated.

| Cases | $Q_{10}$ values | $Q_{10m}$ values |
|---|---|---|
| $Q_{10} > Q_{10m}$ | 1.88 | 1.5 |
| $Q_{10} = Q_{10m}$ | 1.88 | 1.88 |
| $Q_{10} < Q_{10m}$ | 1.5 | 1.88 |
| Fish | 1.88 | 1.88 |

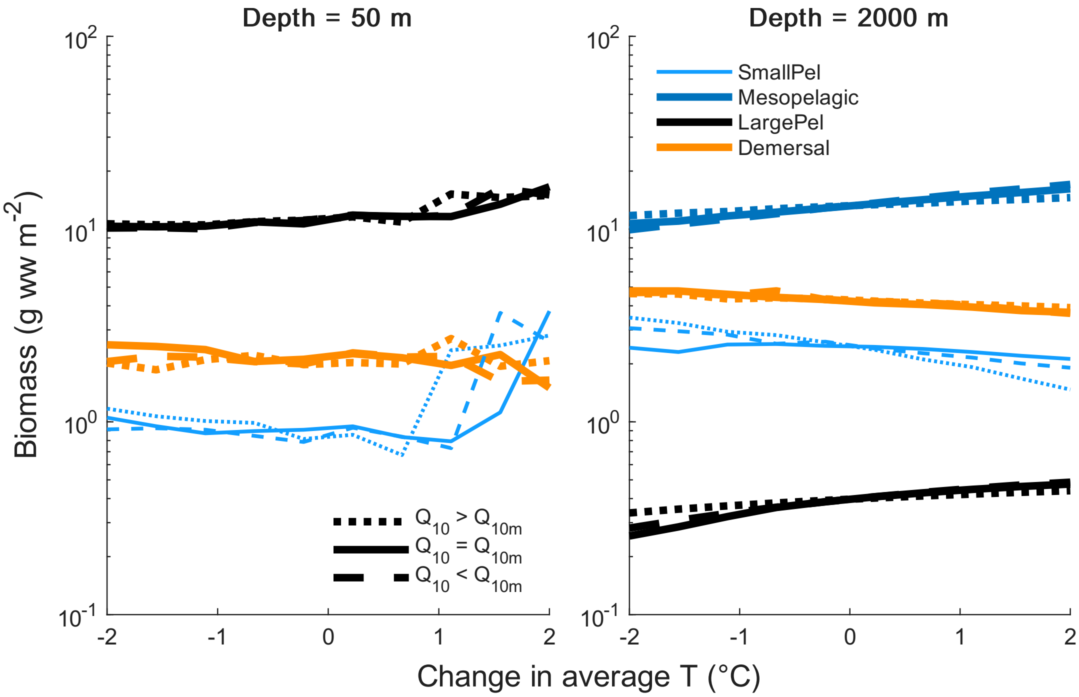


Figure SB1: Simulating an average increase and decrease of temperature of ±2 °C in FEISTY-Squid. Simulations are made for the three $Q_{10}$ cases for the two types of systems.

### Supplement C: Effect of varying $Q10$ assumptions between squid and fish.

Here we tested varying assumptions for the temperature sensitivity of physiological rates ($Q_{10}$) between fish and squid. Unlike in Supplement B, we assumed $Q_{10} = Q_{10m}$ for both groups. In case 1, $Q_{10,\text{ Fish}}$ exceeds $Q_{10,\text{ Squid}}$; in case 2, they are equal; in case 3, $Q_{10,\text{ Fish}}$ is lower than $Q_{10,\text{ Squid}}$ (Table SC1). We kept fish $Q_{10,\text{ Fish}} = 1.88$ and varied $Q_{10,\text{ Squid}}$ from 1 to 2.5, representing extreme values given the limited data on squid $Q_{10}$ (see Discussion). Across all cases, squid biomass decreased with increasing temperature (Fig. SC1), and the choice of $Q_{10}$ did not alter this pattern.

Table SC1: Cases of $Q_{10,\text{ Fish}}$ values for squid tested for temperature simulations. $Q_{10,\text{ Fish}}$ values of fish are also indicated.

| Cases | $Q_{10,\text{ Fish}}$ values | $Q_{10,\text{ Squid}}$ values |
|---|---|---|
| $Q_{10,\text{ Fish}} > Q_{10,\text{ Squid}}$ | 1.88 | 1 |
| $Q_{10,\text{ Fish}} = Q_{10,\text{ Squid}}$ | 1.88 | 1.88 |
| $Q_{10,\text{ Fish}} < Q_{10,\text{ Squid}}$ | 1.88 | 2.5 |

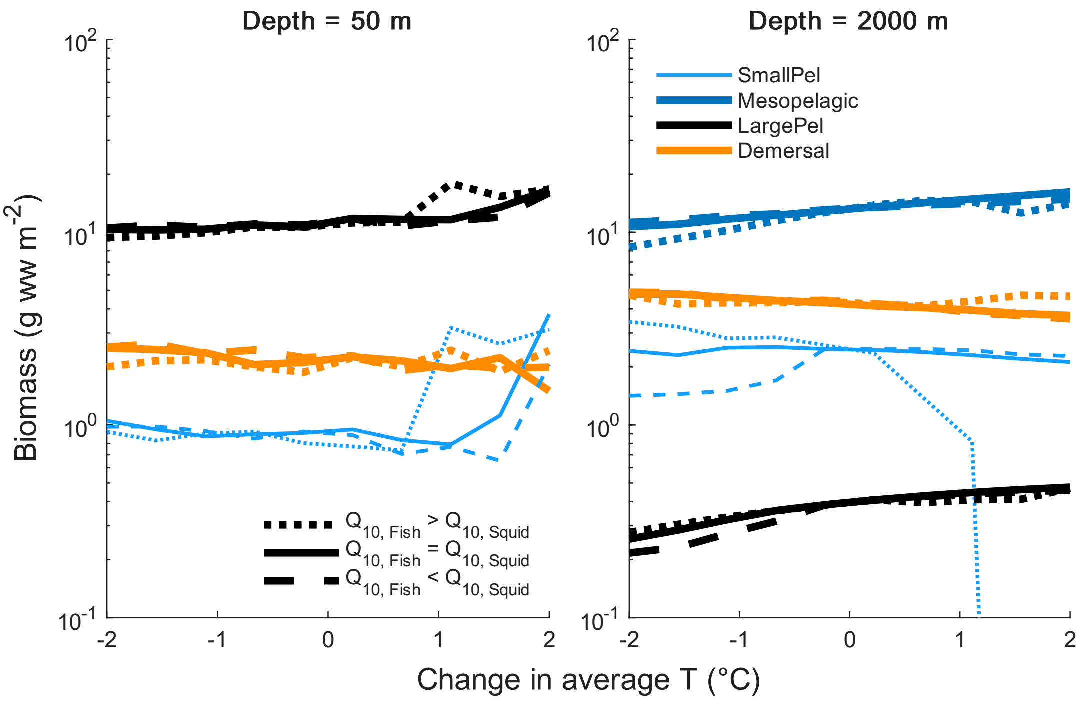


Figure SC1: Simulating an average increase and decrease of temperature of ±2 °C in FEISTY-Squid. Simulations are made for the three $Q_{10}$ cases for the two types of systems.
